## Supplementary figures and images for "The timing of transcription of RpoS-dependent genes varies across multiple stresses in *Escherichia coli* K-12"

### Figure S1

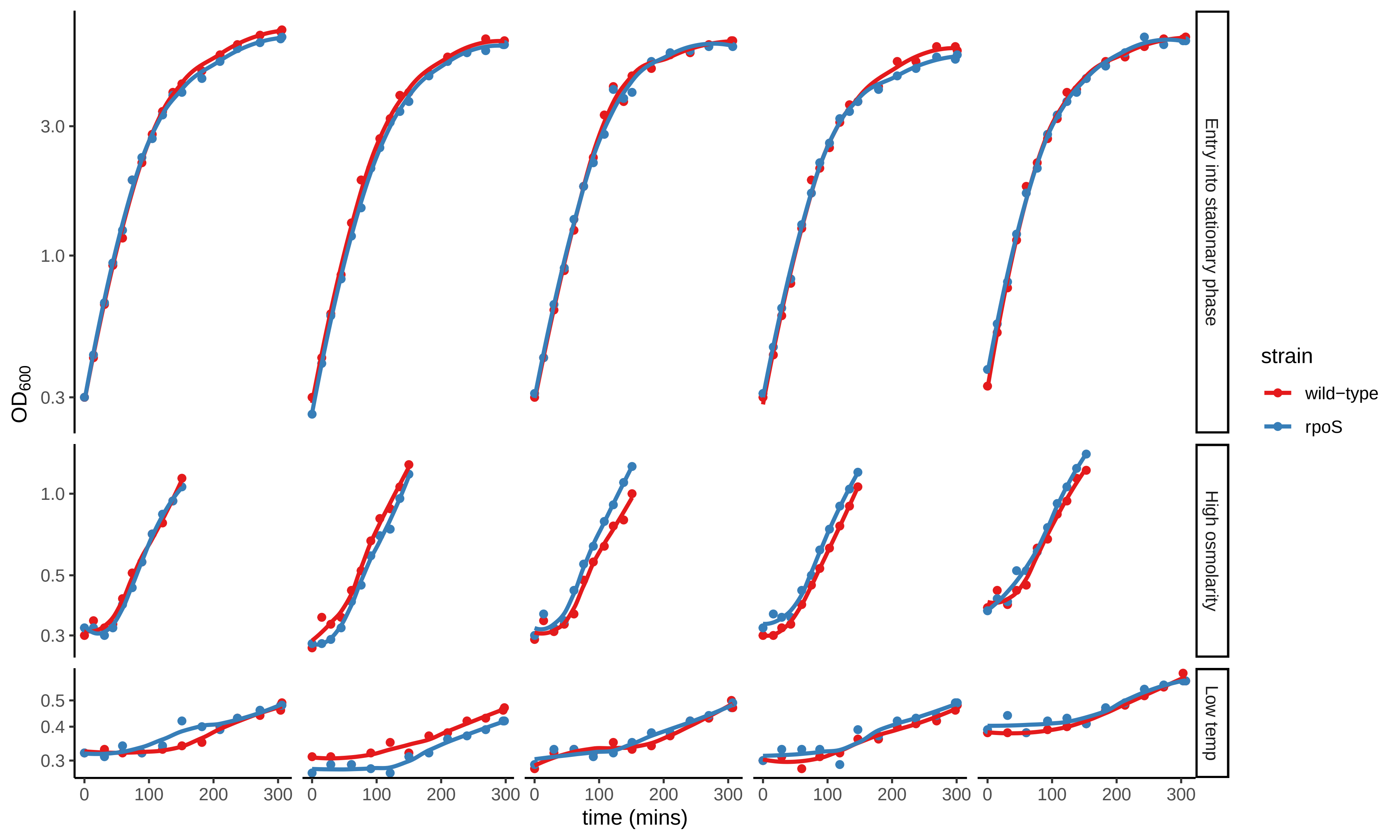
