## Supplementary material for "The timing of transcription of RpoS-dependent genes varies across multiple stresses in *Escherichia coli* K-12": Figure S2

A

Stationary Phase

| Lane | 9 | 10 | 11 | 12 | 13 | 14 | 15 |
|---|---|---|---|---|---|---|---|
| OD | 0.27 | 0.55 | 1.06 | 1.90 | 2.60 | 3.60 | Δrpos |
| Time | 0 min | 30 min | 60 min | 90 min | 120 min | 150 min | 150 min |

RpoS

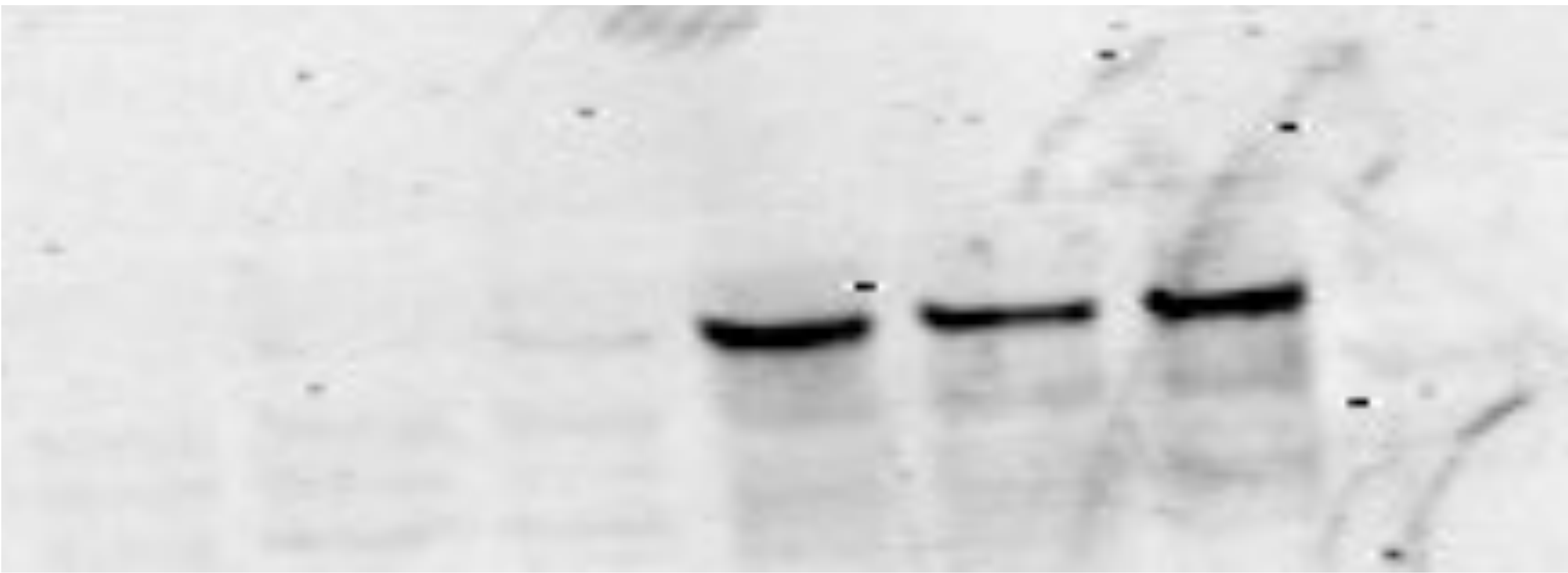

B

High Osmolarity

| | | | | | | |
|---|---|---|---|---|---|---|
| 9 | 10 | 11 | 12 | 13 | 14 | 15 |
| 0.58 | 0.5 | 0.5 | 0.8 | 1.2 | 1.3 | Δrpos |
| 0 min | 30 min | 60 min | 90 min | 120 min | 150 min | 150 min |

RpoS

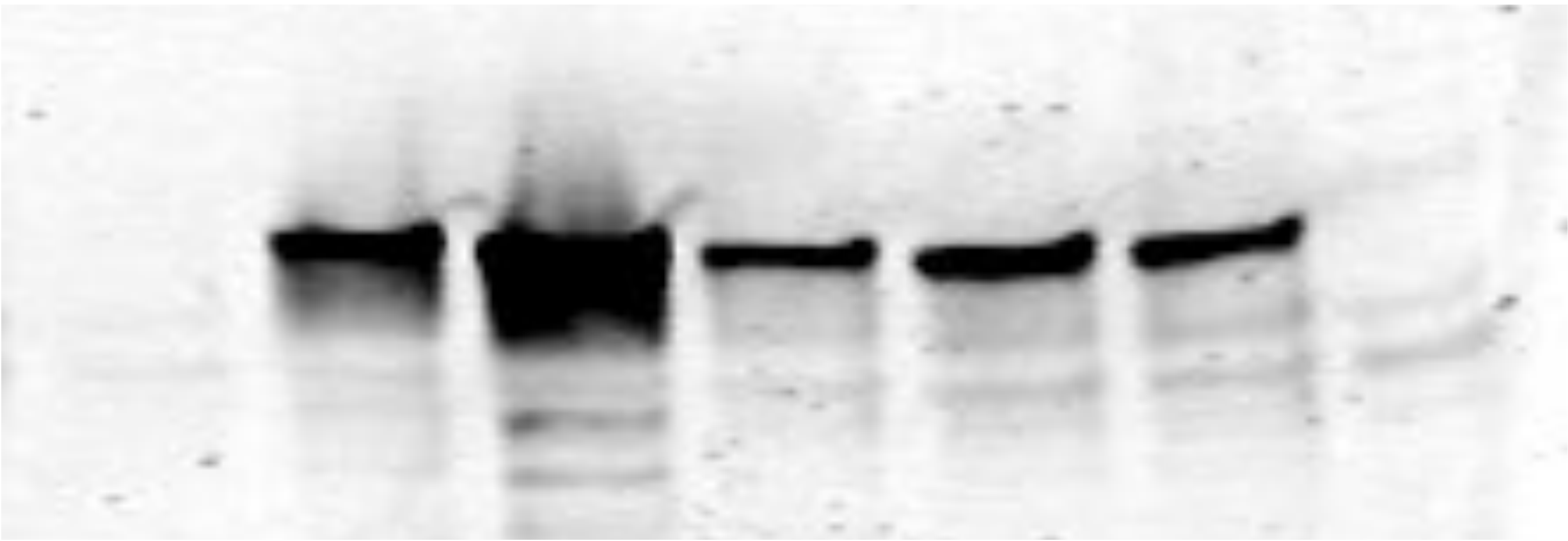

C

Low Temperature

| Lane | 1 | 2 | 3 | 4 | 5 | 6 | 7 | 8 | 9 | 10 | 11 | 12 | 13 |
|---|---|---|---|---|---|---|---|---|---|---|---|---|---|
| OD | Ladder | 0.32 | 0.35 | 0.36 | 0.35 | 0.37 | 0.39 | 0.41 | 0.45 | 0.46 | 0.48 | 0.52 | 0.49 |
| Time since cold shock | | 0 | 29 | 63 | 92 | 121 | 153 | 183 | 220 | 244 | 274 | 301 | Δrpos |

RpoS

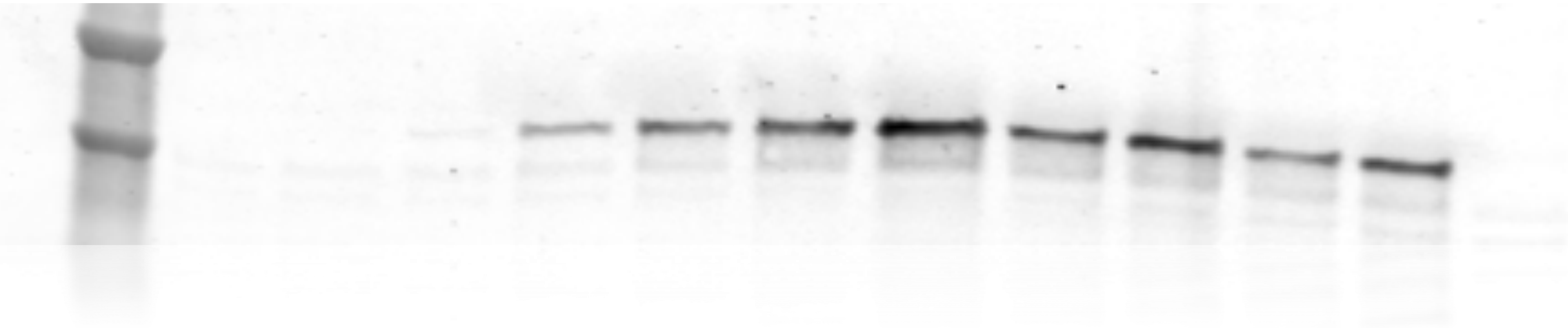
